# Metformin prevented high glucose-induced endothelial reactive oxygen species via OGG1 in an AMPKα-Lin-28 dependent pathway

**DOI:** 10.1101/2020.11.18.388462

**Authors:** Liangliang Tao, Xiucai Fan, Jing Sun, Zhu zhang

## Abstract

Metformin improved vascular function in obese type 2 diabetic patients. 8-oxoguanine glycosylase (OGG1), a main DNA glycosylase, was involved in vascular complications in diverse diseases. However, whether metformin suppressed endothelial ROS via OGG1 pathway was unclear. Human umbilical vein endothelial cells (HUVECs) were exposed to HG (high glucose) or HG with metformin. OGG1 and AMPfα levels were measured after metformin treatment while HG-caused ROS was measured by DHE prober. Diabetic mice were induced by daily intraperitoneal injections of streptozotocin (STZ). Metformin reduced Endothelial ROS caused by HG via upregulating OGG1. Additionally, OGG1 protein expression was dependent on its mRNA stability, which was reversed by genetic inhibition of AMPKα and Lin-28. The role of OGG1 on ROS stimulated by HG was partially dependent on NFKB/NOX4 pathway in HUVECs. These results suggested that metformin contacted HG-induced endothelial ROS via AMPKα/Lin-28/OGG1 pathway.

## Introduction

Vascular complications of Diabetics severely hampered the quality of life and life expectancy of diabetic patients via causing the end-stage renal disease and blindness, and so on[1, 2]. Endothelial reactive oxygen species (ROS) induced by hyperglycemia is the pathological basis of vascular complications and the potential clinical treatment target[3, 4].

AMP-activated protein kinase (AMPK), consisting of a catalytic α subunit and regulatory β and γ subunits, is the master regulator of energy metabolism[5, 6]. It was activated in response to energy stress such as low glucose, hypoxia, stroke and ischemia and so on[7, 8]. Metformin was widely used to control the glucose level in the diabetic patients via AMPK activation. Metformin inhibited mortality in myocardial infarction and heart failure patients, the morbidity of cardiovascular events in heart failure and type II diabetes mellitus patients[9], and the incidence and severity of stroke in patients with type 2 diabetes mellitus[10]. The underlying mechanism of Metformin-mediated vascular function attracted our attention.

8-oxoguanine glycosylase-1 (OGG1) is one of DNA glycosylase enzymes for the repair of the DNA Lesion8-oxoguanine (8-oxoG) induced by oxidative stress[11]. Emerging evidence showed the OGG1 was involved in oxidative stress-related pathological progress[12, 13]. OGG1 deficiency exhibited lower levels of DNA strand lesions, PARP1 overactivation and promoted cell death induced by hydrogen peroxide, suggesting the role of OGG1 on genome integrity[12]. Gene deletion of OGG1 promoted cells resistant to oxidative stress-induced DNA demethylation caused by oxidative stress[13]. However, the role of OGG1 on endothelial function in type II diabetes mellitus patients was still unclear.

In this study, we found that AMPK inhibitor metformin remarkably prevented endothelial oxidative stress via upregulating OGG1 in response to high glucose.

## Materials and methods

### Animal models

The protocols for the animal experiments were approved by the Institutional Animal Care and Use Committee of Nanjing Medical University (Nanjing, China). The 8-week-old male C57/BL6 mice weighting 22-25g were injected intraperitoneally with streptozotocin (STZ, 55 mg/kg) for 5days[14]. 7 days later, mice with blood glucose ≥16.7 mmol/L were diagnosed with diabetes, and then sacrificed and the aortas were dissected and embedded into the OCT liquid.

### Cell culture

The human umbilical vein endothelial cell (HUVEC) line EA.hy926 was obtained from Cell Bank of the Chinese Academy of Sciences, maintained in a humidified incubator at 37°C with 5% CO^2^. HUVECs were exposed to high glucose (30 mM) or metformin with high glucose for indicated different time periods. Metformin was bought from Selleck company (S1950).

### Western blotting

Protein samples were obtained from HUVECs preconditioned with HG. Bradford (BCA) assay was used to quantify the concentration of above samples. Cell lysates were subjected to Western blotting with specific antibodies. Primary antibodies anti-OGG1 (NB100-106, Novus Biologicals), anti-Lin28 ser 200 (20607, CST), anti-Lin28 (3695, CST), anti-AMPK (5831, CST), anti-Phospho-AMPKα (50081, CST), and β-Actin (4970, CST) were obtained commercially.

### Immunofluorescence

Briefly, tissue slides were fixed in 4% paraformaldehyde at room temperature (RT) for 15 min, following wash in PBS, and permeabilization with 0.2% Triton X-100 in PBS for 10 min at RT. Then the slides were cultured with the primary antibodies (dilution 1:200) for overnight at 4°C after blocking with 5% BSA for 30 minutes. DAPI was used for nuclear stain (dilution 1:100; Beyotime). Laser scanning confocal microscopy (LSM 710; Carl Zeiss, Germany) was used to acquire the images.

### Quantitative real-time PCR

Total RNA was extracted with Trizol reagent, as described before[15]. Then we measured the concentration of RNA via NanoDrop 2000 (Thermo Scientific). mRNA (1 μg) was converted to cDNA, and then cDNA was amplified using following the primers listed below. 18S was used as a control. 18S F TGTGCCGCTAGAGGTGAAATT, R TGGCAAATGCTTTCGCTTT OGG1 F CACACTGGAGTGGTGTACTAG, R CCAGGGTAACATCTAGCTGGAA.

### Small interfering RNA and overexpression experiments

After growing to approximately 80% confluency in the 6-well plates, the cells were transfected with 8μl (10 μM) small interfering RNA (sc-43983, Sant Cruz Biotechnology) with 8 μl Lipofectamine™ RNAiMAX (Invitrogen) for 6 hours according the manufacturer’s instructions. After culturing for 24 hours, the cells were exposed for HG for another indicated time.

Cells in the 6-well plates were transfected with 2μg plasmid of AMPKα or OGG1 via 6 μl Lipofectamine 2000 (Invitrogen) for 6 hours following the manufacturer’s instructions. After appropriated incubation, the cells were lysed for western blotting.

### ROS and measurements

According to the manuscript’s protocol, HUVECs were incubated with DHE fluorescent dye (50 μM, Beyotime Institute of Biotechnology, China) at 37°C for 30 min after cocultivation with HG or not[16]. Fluorescence microscopy was applied to photograph the cells.

### Statistical results

All data are showed as the means ± standard error of the mean. Data were analyzed by using Student’s t test or one-way analysis where it was appropriate. Values of P< 0.05 were considered statistically significant. All statistical analyses were performed using GraphPad PRISM software 5.0 (GraphPad Inc., USA). Bar charts represent the mean values ± standard error of the mean of at least three independent trials.

## Results

### High glucose decreased OGG1 expression and increased endothelial reactive oxygen species

At first, we measured the OGG1 protein and mRNA expression after high glucose (HG) exposure for 0, 6, 12 and 24 hours. Consistent with previously studies[15, 17], our results showed that HG significantly reduced OGG1 expression (Fig.1A-C). Similarly, STZ-induced diabetic mice showed OGG1 level was decreased in the endothelium of aorta compared to control mice (Fig. 1D). Additionally, we also investigated the endothelial reactive oxygen species (ROS) after high glucose treatment for 24 hours via fluorescent probe-DHE (10 μM). The results showed high glucose increased endothelial ROS levels (Fig. 1E-F).

**Figure 1.**
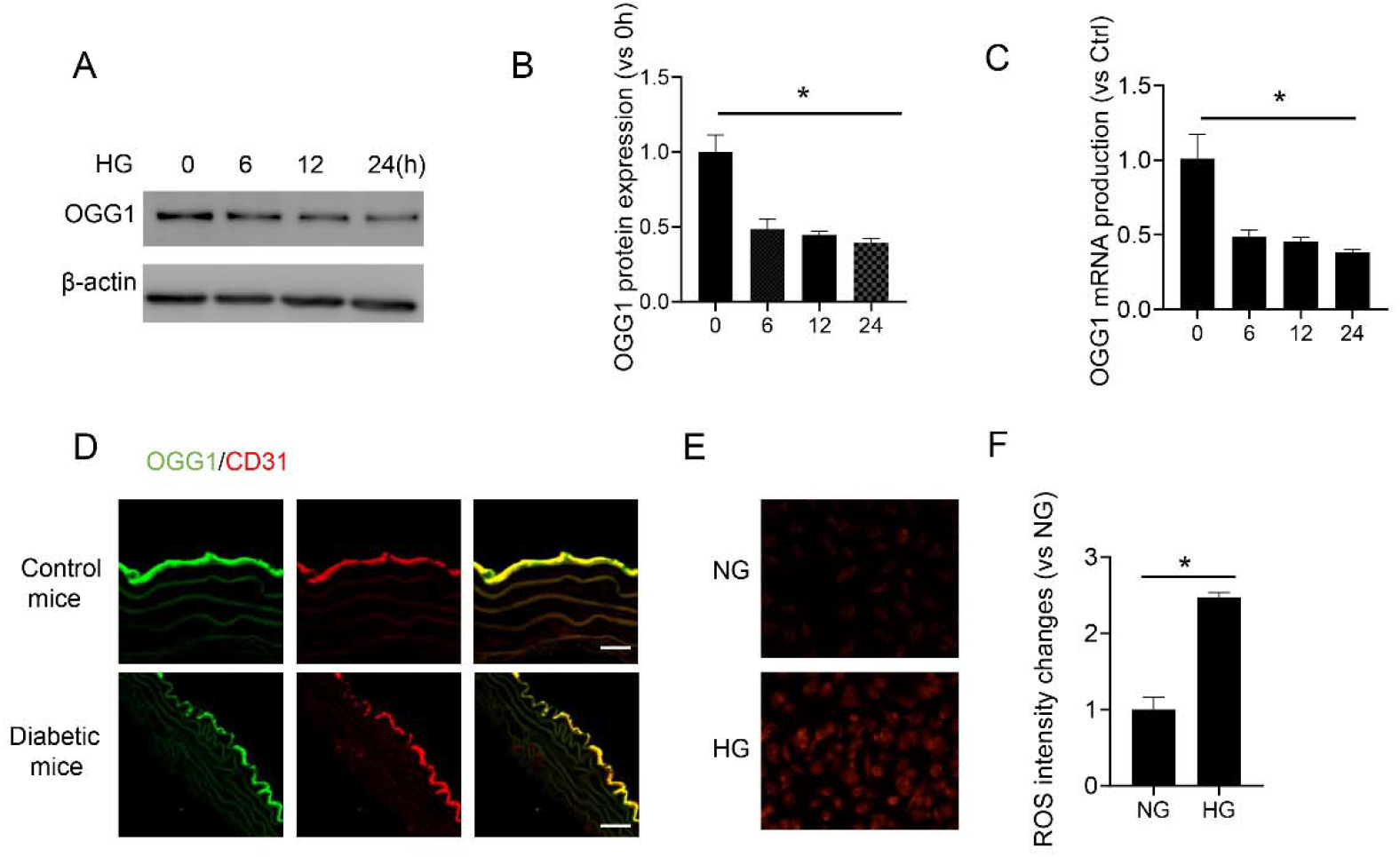
High glucose decreased OGG1 expression while increased endothelial reactive oxygen species levels. (A-C) OGG1 mRNA and protein levels were tested after HG exposure for indicated periods. (B) is the statistical result of (A). (D) The expression of OGG1 in the aorta of diabetic mice were assessed via Immunofluorescence. (E-F) HUVECs were digested and incubated with DHE probe (1 μM) for 30 minutes after high glucose exposure for 12 hours. The mean intensities were obtained via flow Jo software. Scar bar =25μm. *P<0.05 versus NG group (mannitol control group). Values are the means±SD with N = 3.

### Metformin decreased endothelial ROS via OGG1 expression

Metformin was a known and widely used anti-diabetic agent. However, whether the Metformin improved vascular function of diabetic patients via OGG1 was unknown. At first, endothelial ROS was measured after high glucose incubation with metformin or not. The results showed that metformin significantly decreased endothelial ROS (Figure. 2A-B). Then we tested the OGG1 protein levels and mRNA with metformin (0.5 mM) incubation for 24 hours. Metformin increased OGG1 mRNA and protein levels (Fig. 2C-E).

**Figure 2.**
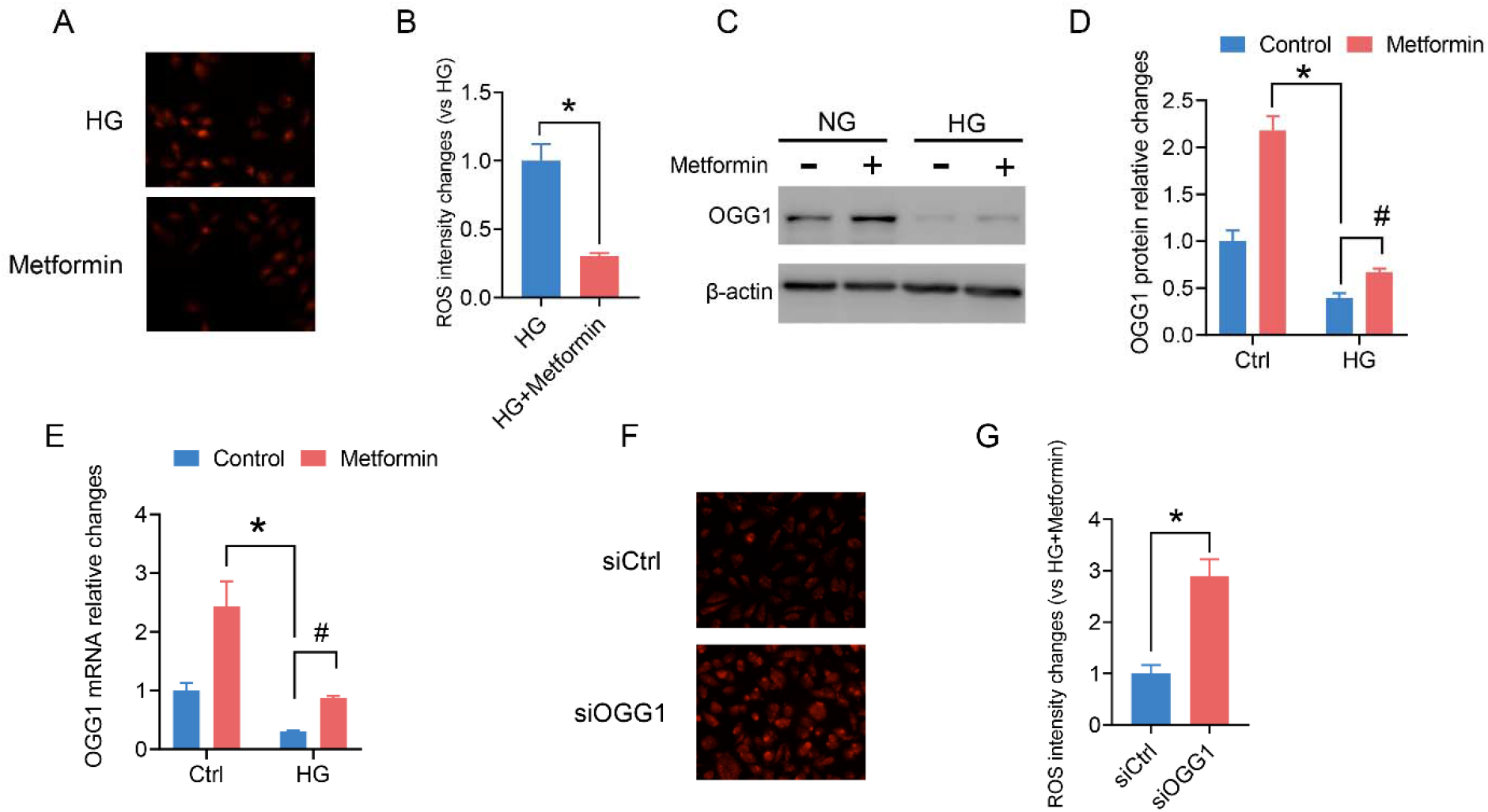
Metformin decreased ROS and activated OGG1 in the HUVECs. (A-B) Endothelial ROS was measured by DHE probe after high glucose exposure with metformin or not. (B) is the stastical result of (A). (C-E) OGG1 mRNA and protein levels were tested after metformin (0.5 mM) exposure for 24 hours. (E) is the statistical result of (D). *P<0.05 versus NG group (mannitol control group). #P<0.05 versus HG group. Values are the means±SD with N = 3. (F-G) HUVECs were digested and incubated with DHE probe (1 μM) for 30 minutes after silencing OGG1 or not in the presence with HG and metformin. The mean intensities were obtained via flow Jo software. Scar bar =25μm. *P<0.05 versus HG plus metformin group. Values are the means±SD with N = 3.

OGG1 was reported to involve in the endothelia ROS production by HG. Herein, we measured the role of metformin in ROS levels after silencing OGG1. Without exception, OGG1 deletion significantly blocked the inhibitory effect of metformin on HG-induced endothelial ROS (Fig. 2F-G), suggesting metformin decreased endothelial ROS via OGG1 production.

### Metformin increased OGG1 protein levels via mRNA stability in response to high glucose

Subsequently, the underlying mechanism of OGG1 upregulation induced by metformin attracted our attention. Thus, we performed mRNA degradation assays using actinomycin D (Act D, 10 μg/ml) to inhibit de novo mRNA transcription. qPCR analysis showed OGG1 mRNA half-life time was significantly increased in response to metformin, suggesting metformin mainly increased OGG1 mRNA stability (Fig. 3A). Additionally, we tested protein degradation assays using cycloheximide (CHX, 10 μg/ml)) to prevent protein translation. Western blot exhibited that metformin did not suppress endothelial OGG1 expression (Fig. 3B-C). All results exhibited that OGG1 protein levels by metformin were dependent on its mRNA stability.

**Figure 3.**
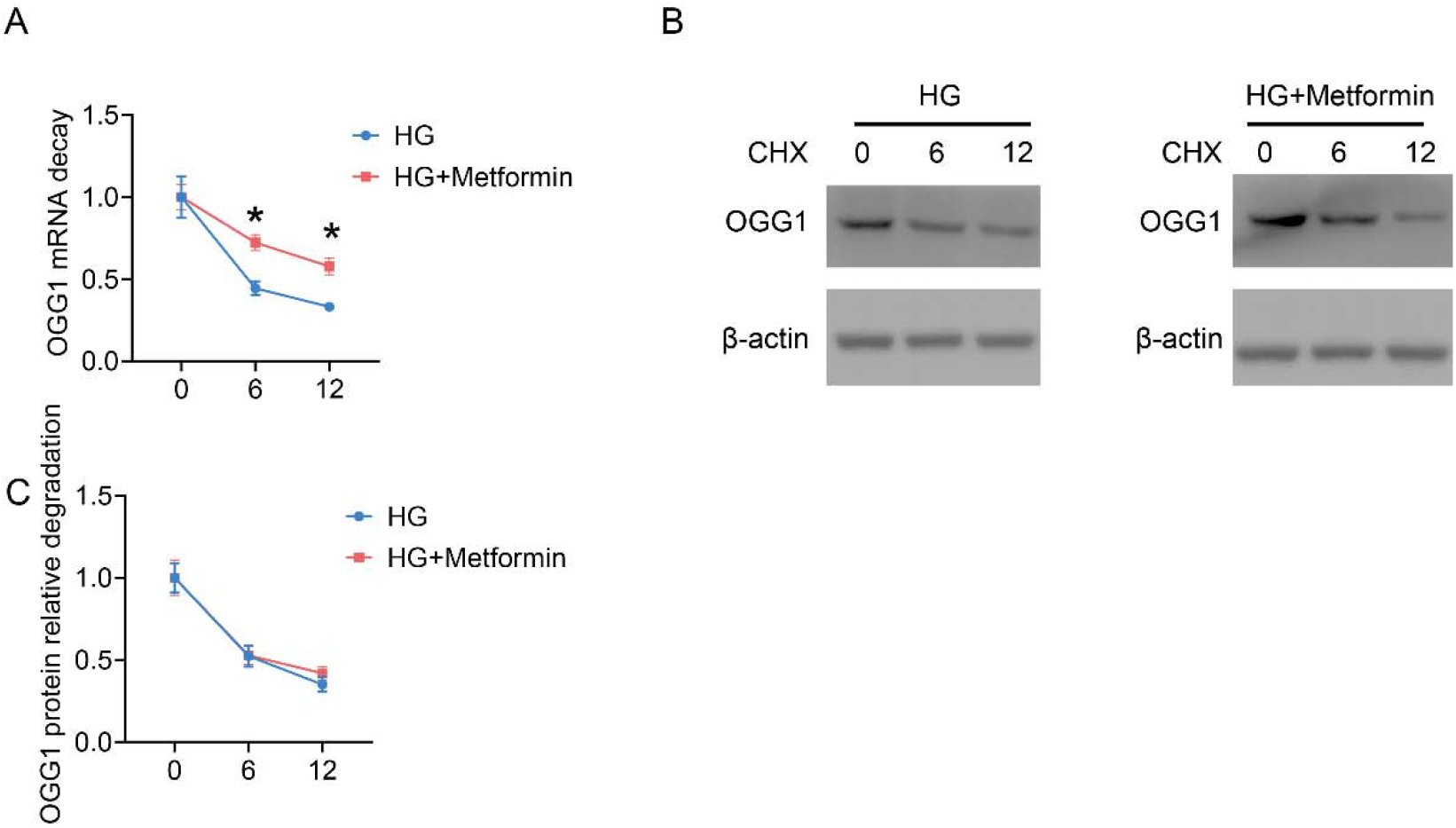
The OGG1 expression stimulated by metformin was dependent on its mRNA stability in the HUVECs. (A) HUVECs were incubated with actinomycin D (Act D, 10 μg/ml) for 0, 6, and 12 hours, and then OGG1 mRNA was measured by qPCR. (B-C) OGG1 protein levels were tested by WB after CHX-treated HUVECs for 0, 6, and 12 hours. *P<0.05 versus control group (mannitol control group). Values are the means±SD with N = 3.

### AMPKα-mediated OGG1 expression was dependent on its mRNA stability

Concerning that AMPKα is a key downstream target of metformin and reported to regulate mRNA stability[18], we investigated whether OGG1 upregulation by metformin was dependent on AMPKα activation. Metformin significantly upregulated AMPKα in the HUVECs (Fig. 4A-B), while silence of AMPKα abrogated metformin-induced OGG1 expression (Fig. 4C-E), suggesting AMPKα regulated OGG1 level through increasing its mRNA stability. Additionally, we also explored the role of AMPKα on endothelial ROS. It was clear that silence of AMPKα predominantly upregulated endothelial ROS levels (Fig. 4F-G)

**Figure 4.**
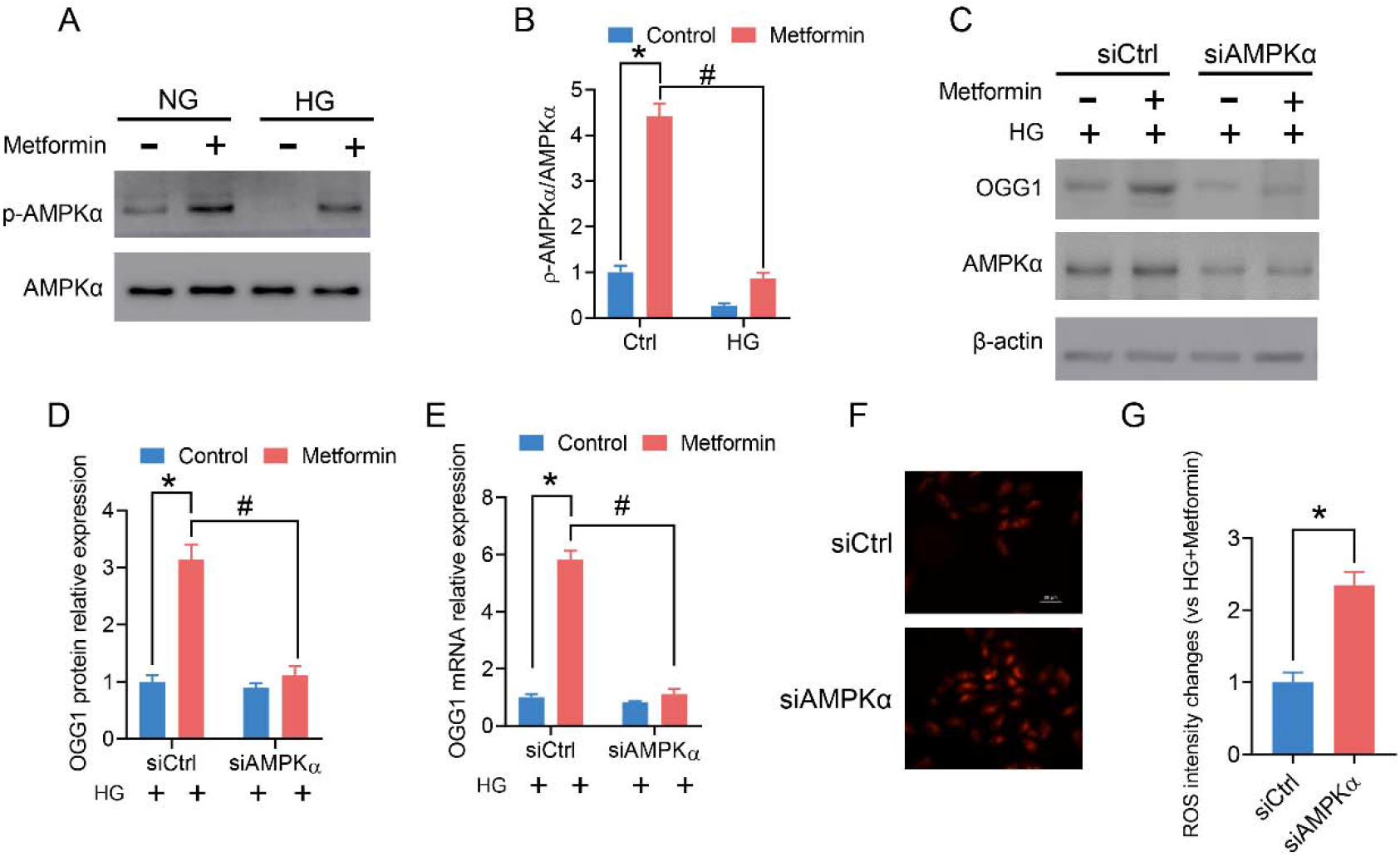
The mRNA stability of OGG1 triggered by metformin was dependent on AMPKα. (A-B) The phosphorylation of AMPKα and total AMPKα were measured after metformin-treated HUVECs. (C-E) OGG1 protein and mRNA levels were tested by WB and qPCR after metformin and high glucose treatment simultaneously with silencing of AMPKα or not in the HUVECs. (F-G) HUVECs were digested and incubated with DHE probe (1 μM) for 30 minutes after silencing AMPKα or not in the presence with HG and metformin. The mean intensities were obtained via flow Jo software. Scar bar =25μm. *P<0.05 versus control group. #P<0.05 versus HG plus metformin group. Values are the means±SD with N = 3.

### Lin28 stabilized OGG1 mRNA levels after high glucose treatment

To further identify the role of AMPKα in OGG1 regulation by metformin, we assessed RNA binding proteins (RBPs), which play significant roles in the regulation of mRNA stability[19]. Lin-28 is a member of RNA-binding proteins, and one potential target of AMPKα [20], Additionally, the role of Lin-28 mainly depended on its cytoplasmic sublocation. We test the Lin-28 cytoplasmic distribution after metformin exposure in the presence of HG or not. The results showed metformin promoted HG-suppressed Lin-28 cytoplasmic distribution (Fig. 5A-B). Subsequently, silencing Lin-28 significantly reduced OGG1 mRNA and protein levels induced by metformin, showing that Lin-28 was the key upstream of OGG1 mRNA stability (Fig. 5C-E). Additionally, the OGG1 mRNA stability also measured after gene deletion of Lin-28 in the presence of actinomycin D (Act D, 10 μg/ml). Silence of Lin-28 significantly decreased OGG1 mRNA stability (Fig. 5F)

**Figure 5.**
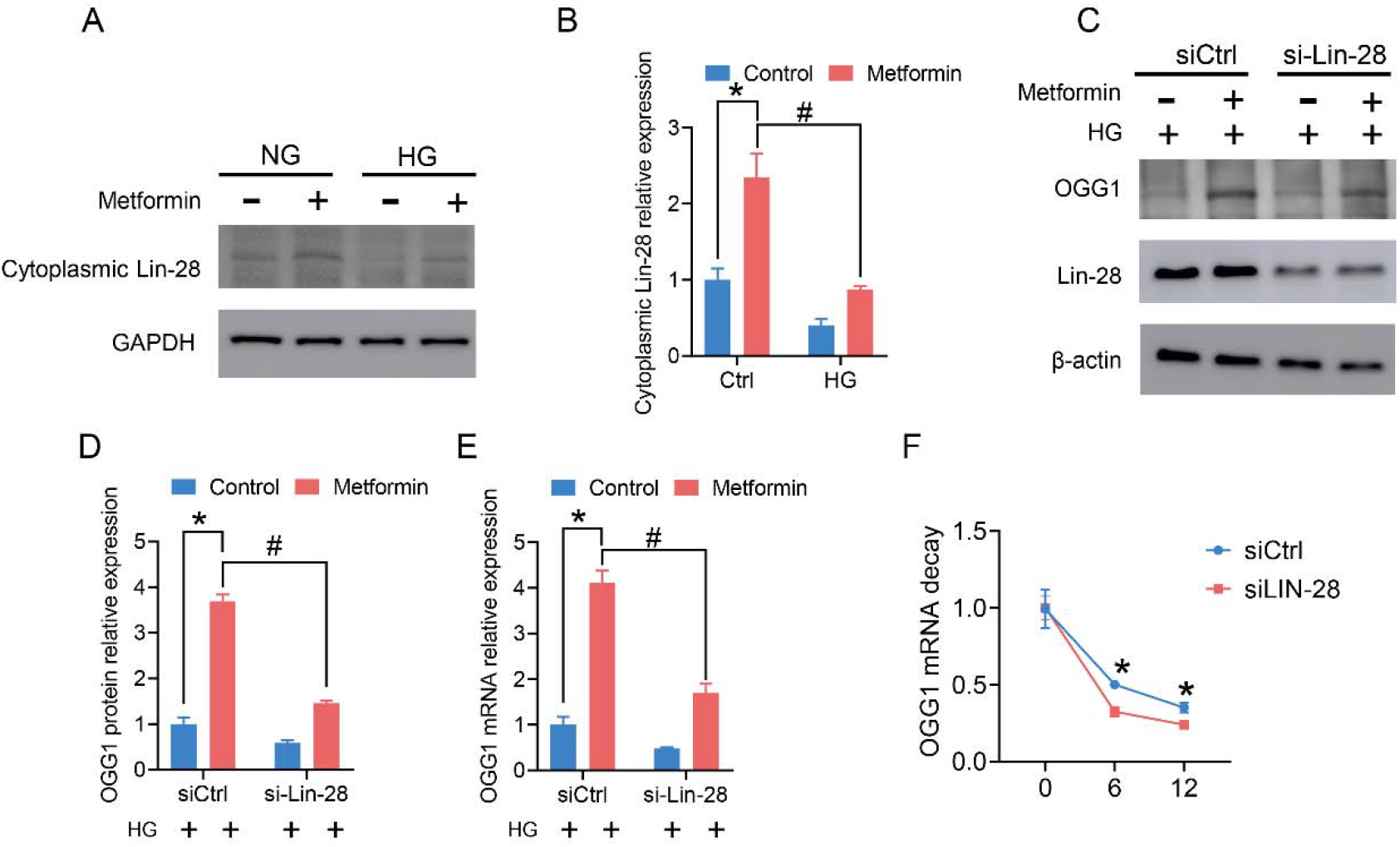
Lin28 meditated OGG mRNA stability of OGG1 triggered by metformin. (A-B) The subcellular localization of Lin28 were measured after metformin-treated HUVECs in the presence of high glucose. (C-E) OGG1 protein and mRNA levels were tested by WB and qPCR after metformin and high glucose treatment simultaneously with silencing of Lin28 or not in the HUVECs. (F) The half-time hours of OGG1 mRNA was measured after silencing Lin-28 or not with actinomycin D (Act D, 10 μg/ml). *P<0.05 versus control group. Values are the means±SD with N = 3.

### Metformin activated OGG1 via Lin-28 phosphorylation by AMPKα

Next, we explore the mechanism of metformin on Lin-28 cytoplasmic retention. Since AMPKα was reported to mediate Lin-28 nuclear export through phosphating Lin-28 [20]. According, we speculated metformin induced a dramatic accumulation of Lin-28 in the cytoplasm via AMPKα activity. Silence of AMPKα or AMPKα inhibitor with Compound C (10 μM) attenuated the Lin-28 cytosolic location and phosphorylation (Fig. 6A-B), suggesting the role of AMPKα in the Lin-28 distribution and activity. Further rescue experiments indicated that silencing Lin28 significantly reduced AMPKα-induced OGG1 expression induced by AMPKα activation (AICAR) or AMPKα plasmid (Fig. 6C-D).

**Figure 6.**
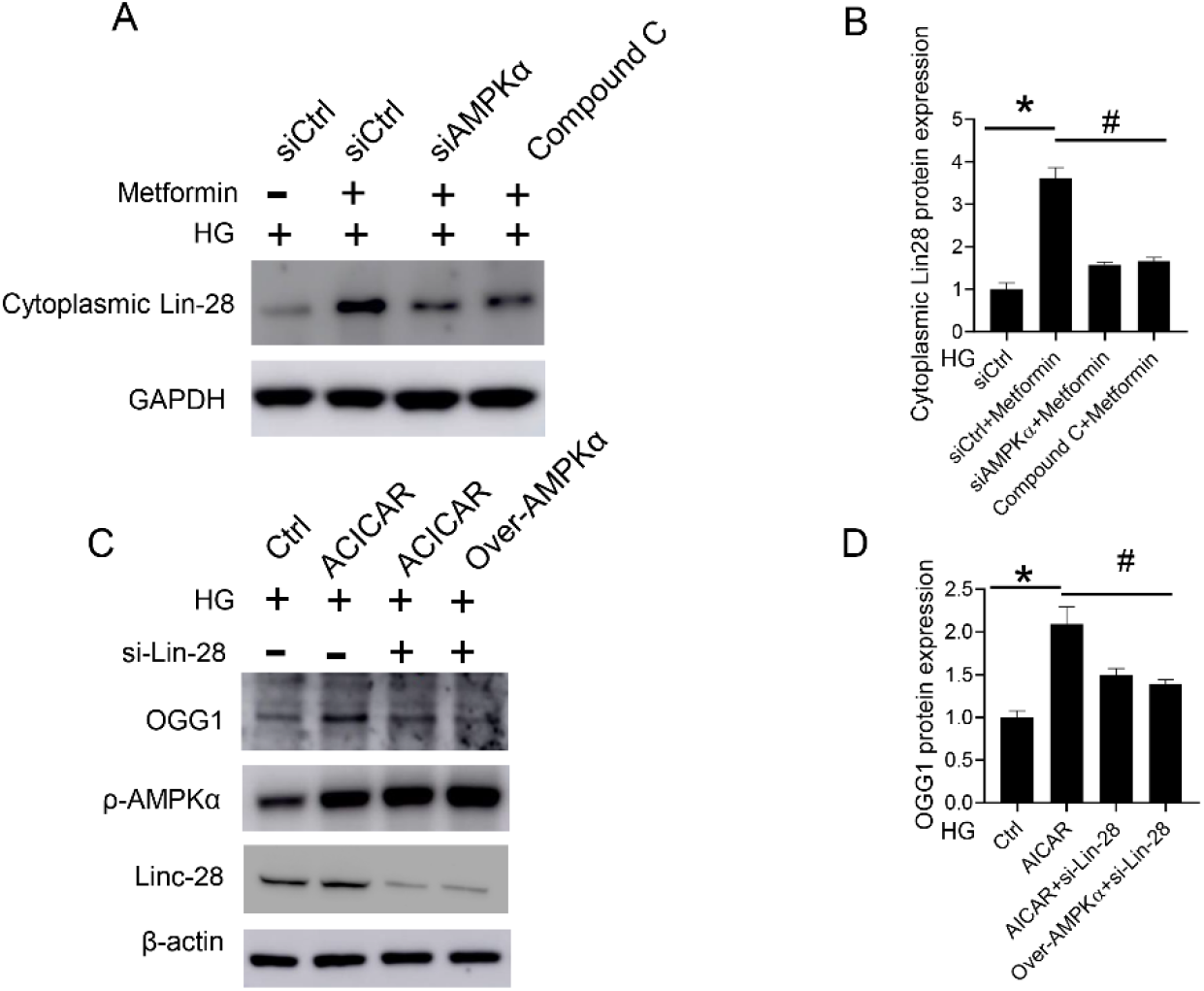
OGG1 expression induced by metformin was dependent on AMPKα phosphorylation of Lin-28. (A-B) The subcellular localization of Lin28 were measured after AMPKα silence or inhibitor pretreatment. *P<0.05 versus HG group. #P<0.05 versus HG plus metformin group. (C-D) OGG1 protein and mRNA levels were tested after silence Lin28 in the HUVECs with AMPKα plasmid or AICAR. *P<0.05 versus HG group. #P<0.05 versus HG plus ACICAR group. Values are the means±SD with N = 3.

### Metformin decreased endothelial reactive oxygen species via AMPKα/Lin-28/OGG1 pathway

Lastly, we assessed the role of AMPKα/Lin-28/OGG1 pathway on metformin-prevented endothelial reactive oxygen species. HUVECs were pretreated with AMPKα inhibitor Compound C or gene deletion of Lin-28. The results exhibited that both Compound C or interference by siRNA of Lin-28 predominantly accelerated endothelial ROS production caused by high glucose (Fig. 7A-B).

**Figure 7.**
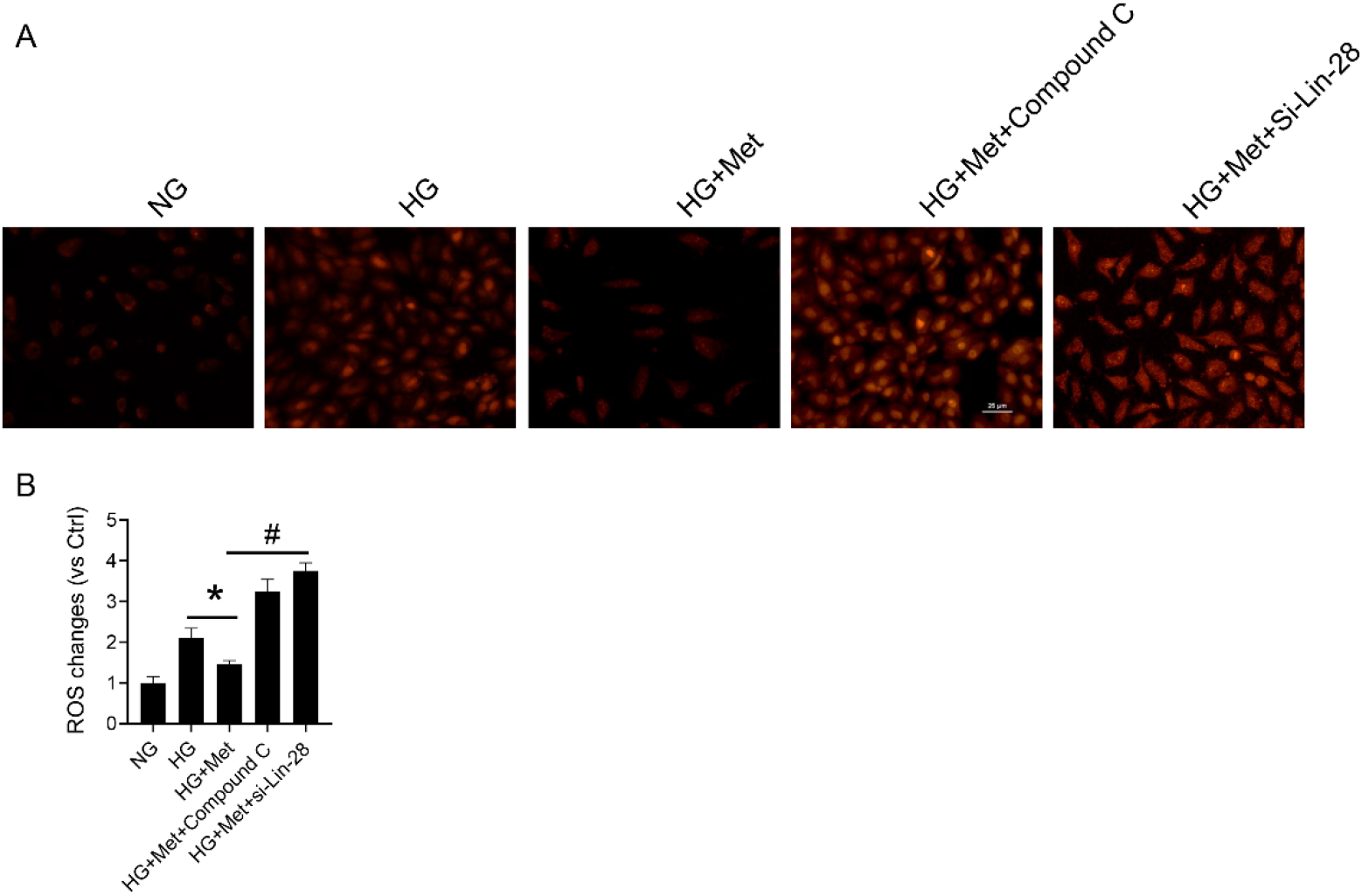
The protective role of metformin on endothelial ROS was dependent on AMPKα/Lin-28/OGG1 pathway. (A-B) Endothelial ROS levels were assessed by DHE after Compound C or silence of Lin-28-treated HUVECs. *P<0.05 versus HG group. #P<0.05 versus HG plus metformin (Met) group. Values are the means±SD with N = 3.

### OGG1 prevented HG-induced endothelial ROS via NFκB/NOX4 pathway

We next searched for the possible mechanism linking OGG1 and ROS. Previous study reported that NADPH oxidases are the main endogenous resource of endothelial ROS[21]. So we measured the levels of NOX1, NOX2, and NOX4 after OGG1 overexpression with high glucose treatment or not. Forcing expression of OGG1 dramatically abrogated HG-induced NOX4 expression rather than NOX1 and NOX2 (Fig. 8A-B). The endothelial mainly proinflammatory transcriptional factor NFκB has been reported involved in 8-oxoG–induced endothelial NOX4 expression[22]. Then we tested whether OGG1-inhibited NOX4 expression was dependent on NFκB. HUVECs were incubated with HG after overexpression of OGG1. It’s clear that OGG1 significantly downregulated NFκB activity (Fig. 8C-D). As expected, silence of NFκB partially reversed HG induction of NOX4 expression (Fig. 8E-F). Collectively, we demonstrated that OGG1 reduced HG induction of ROS via NFκB/NOX4 pathway.

**Figure 8.**
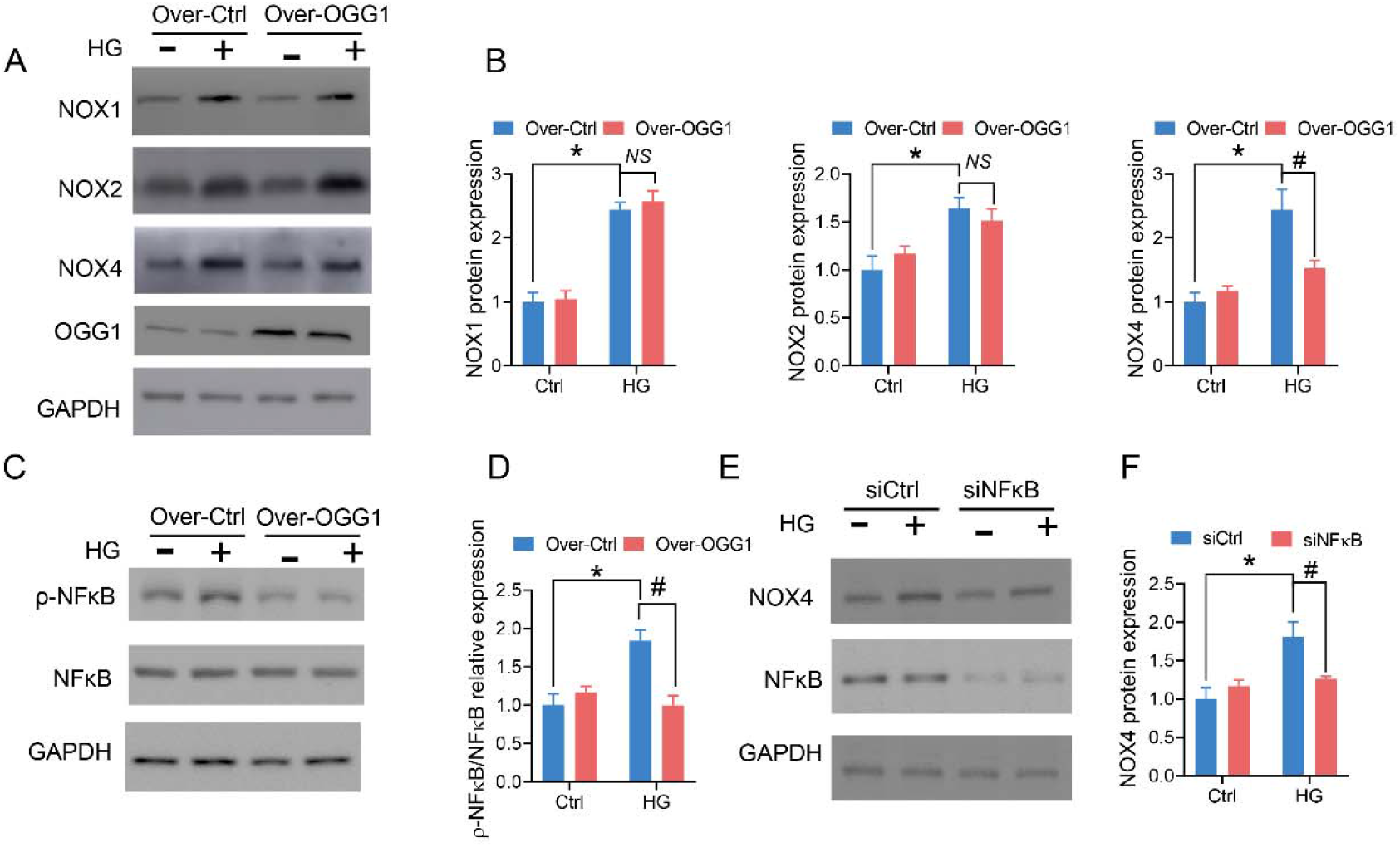
OGG1 regulated HG-induced endothelial ROS via NFκB/NOX4 pathway. (A-B) NADPH oxidases were measured after HUVECs exposure to high glucose treatment with overexpression of OGG1. (B) was statistical result of (A). (C-D) NFκB activity was measured after OGG1 overexpression with high glucose treatment for 24 hours. (E-F) NOX4 protein level was assessed after NFκB silence with high glucose incubation. *P<0.05 versus Ctrl group. #P<0.05 versus HG group. ^*NS*^P means no difference. Values are the means±SD with N = 3.

## Discussion

In this study, we found that metformin protected against endothelial reactive oxygen species via upregulating OGG1. Additionally, the underlying mechanism of metformin in OGG1 expression was dependent on AMPKα/Lin-28 pathway (Fig. 9). Additionally, OGG1 significantly reduced endothelial ROS through NFκB/NOX4 pathway.

**Figure 9.**
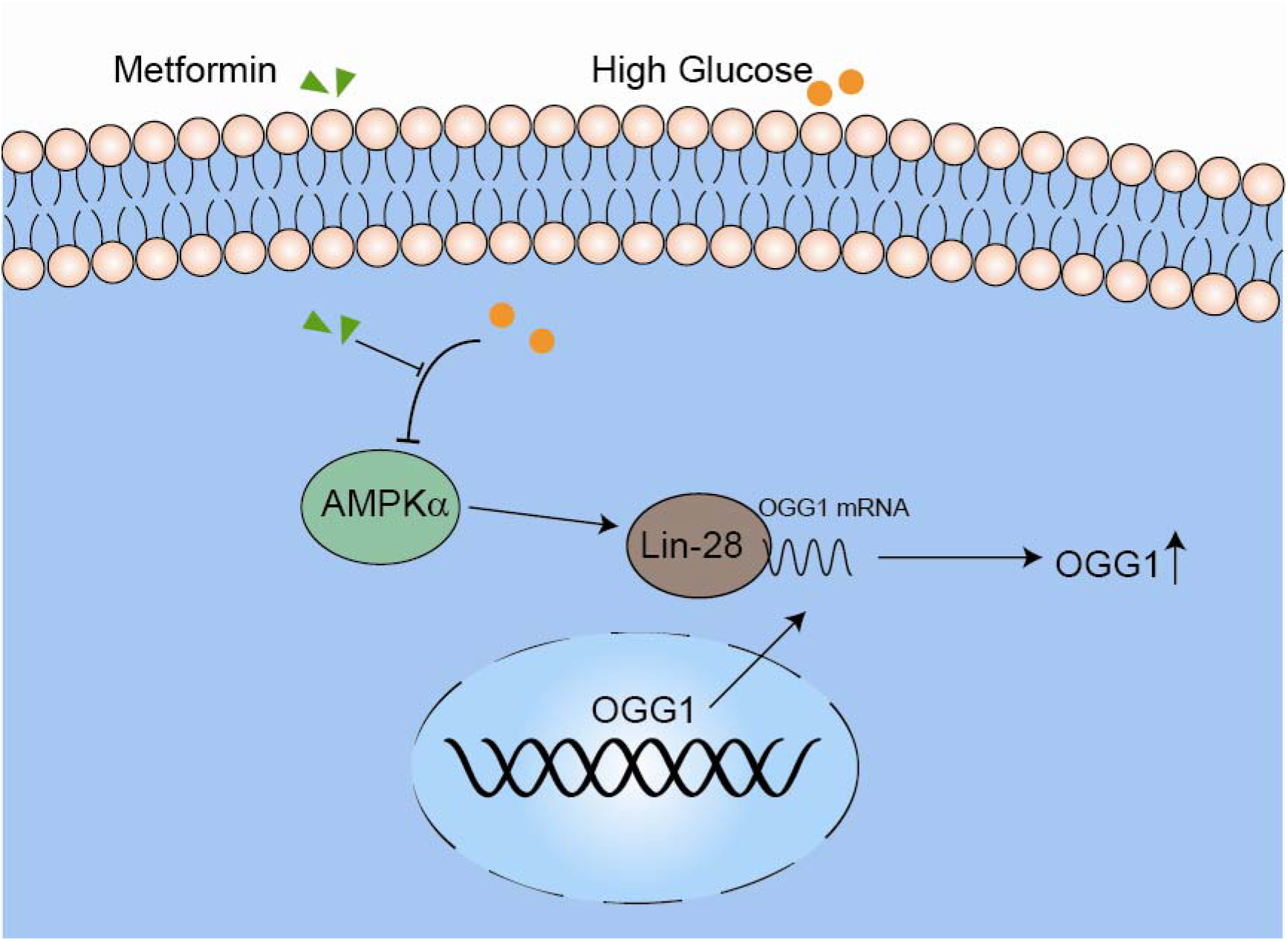
Metformin preserved against high glucose-induced endothelial reactive oxygen species via AMPKα/Lin-28 pathway.

Metformin is the first-line medication for the treatment of diabetes agent, improving the risk of cardiovascular disease among patients with type 2 diabetes, and reducing cardiovascular mortality and morbidity in patients comparison with alternative glucose-lowering drugs[23, 24]. Metformin has also been reported to improve endothelium-dependent vasorelaxation in a NO dependent manner[25, 26]. Additionally, administration of metformin in drinking water displayed an decrease in blood pressure and vascular damage dependently on AMPK/ER stress pathway[27]. Similarly, we also proved the effect of metformin on vascular damage. Metformin incubation predominantly suppressed endothelial ROS induced by high glucose measured by DHE fluorescent dye. Mechanismly, the actions of metformin were dependent on AMP-activated protein kinase (AMPK) or not. AMPK is considered to be a metabolic fuel gauge activated in response to energy depletion[28]. Metformin activated eNOS phosphorylation and NO production in the wild-type but not in AMPK-knockout mice, and overexpression of AMPK-DN (dominant-negative mutation) abrogated the protective effects of metformin on eNOS activity and endothelial apoptosis[29], suggesting the protective role of metformin was dependent on AMPK activity. Further evidence from mouse models with genetic perturbation of AMPKα documented the metformin preserved withdrawal signs precipitated by nicotine withdrawal corroborated dependently AMPKα expression[30]. However, AMPK might be not necessary for the role of metformin on adipogenesis and BMP6-induced hepcidin gene expression[31, 32]. In this study, we assessed whether metformin decreased endothelial ROS in an AMPK pathway. The results exhibited that AMPKα is required for the role of metformin on ROS.

8-Oxoguanine DNA Glycosylase 1 (OGG1), a DNA glycosylase enzyme, was reported to involve in vascular oxidative stress and pathological progression of atherosclerosis[15, 33]. OGG1 expression was controlled by diverse mechanisms. The nuclear transcription factor Y subunit alpha (NF-YA) promoted OGG1 mRNA transcription via binding to its promoter in the presence of TSC-2 gene[34]. Additionally, signal transducer and activator of transcription 1 (STAT1) mediated the transcriptional activity of OGG1 through recruiting and binding to the gamma-interferon activation site (GAS) motif of the OGG1 promoter region in endotoxin-triggered inflammatory response[35]. Furthermore, Rapid and selective inactivation of OGG1 were degraded via binding to CHIP (C-terminus of HSC70-interacting protein) E3 ligase after cells subjection to mild hyperthermia[36]. Surprisingly, we found that OGG1 expression activated by metformin was dependent on mRNA stability after Act D incubation while gene silence of Lin-28 pronouncedly prevented against the OGG1 expression, suggesting the OGG1 levels stimulated by metformin relied on Lin-28 (RNA binding protein). Previously literature showed that AMPKα inhibitor using dorsomorphin promoted dramatic accumulation of Lin-28 in the nucleus and decreased Lin-28 activity. In our study, we also found that AMPKα regulated Lin-28 distribution and activation. However, the mechanism of AMPKα in Lin-28 activity was warranted to explore.

NADPH oxidases are the main resources of reactive oxygen species, consisting of Nox1-5, and the dual oxidases (Duox) 1 and 2 [37]. Among these, NOX1, NOX2 and NOX4 are the isoforms in the endothelial cells. Accumulating evidences corroborate the fundamental function of NOX families on the pathogenesis of diabetes mellitus[38]. Deletion of Nox1or treatment of diabetic ApoE-/- mice with GKT137831 significantly reduced the ability of leukocytes attached to the aortic wall, Macrophage infiltration and Aortic Nitrative and Oxidative Stress compared with diabetic ApoE-/- mice[39]. Additionally, post-hypoxic oxidative stress and programmed cell death in cardiomyocytes exposed to high glucose was attenuated by gene deletion of NOX2[40]. Downregulation of Nox4 by gene deletion of NOX4 in the rat kidney glomerular mesangial cells inhibited the HG-induced augment in mitochondrial superoxide assessed by MitoSOX Red. In this study, we found that OGG1 reduced endothelial reactive species in the HUVECs administrated high glucose through reducing NOX4 expression rather than NOX1 and NOX2. Consistently with previous study[41], NOX4 downregulation by OGG1 was also dependent on NFκB activity. However, the potential mechanism was still unclear, and further research is warranted. Collectively, our results demonstrated that HG induced endothelial ROS, possibly by a mechanism involving in OGG1/NADPH pathway.

In conclusion. Our results showed metformin suppressed high glucose-triggered endothelial reactive oxygen species via OGG1 in an AMPKα-Lin-28 dependent manner.

## Author contributions

Conceptualization and Data curation: Liangliang Tao.

Formal analysis, Methodology and Investigation: Xiucai Fan.

Project administration and Writing - review & editing: Jing Sun.

Funding acquisition and Writing - review & editing: Liangliang Tao and Zhu zhang.

## References

1. Rask-Madsen, C. and G.L. King, Vascular complications of diabetes: mechanisms of injury and protective factors. Cell metabolism, 2013. 17:(1): p. 20–33.

2. Volpe, C.M.O., et al., Cellular death, reactive oxygen species (ROS) and diabetic complications. Cell Death Dis, 2018. 9(2): p. 119.

3. Di Marco, E., et al., Are reactive oxygen species still the basis for diabetic complications? Clin Sci (Lond), 2015. 129(2): p. 199–216.

4. Detaille, D., et al., Metformin Prevents High-Glucose–Induced Endothelial Cell Death Through a Mitochondrial Permeability Transition-Dependent Process. Diabetes, 2005. 54(7): p. 2179–2187.

5. Hardie, D.G., B.E. Schaffer, and A. Brunet, AMPK: An Energy-Sensing Pathway with Multiple Inputs and Outputs. Trends Cell Biol, 2016. 26(3): p. 190–201.

6. Garcia, D. and R.J. Shaw, AMPK: Mechanisms of Cellular Energy Sensing and Restoration of Metabolic Balance. Mol Cell, 2017. 66(6): p. 789–800.

7. Jiang, S., et al., AMPK: Potential Therapeutic Target for Ischemic Stroke. Theranostics, 2018. 8(16): p. 4535–4551.

8. Lin, S.C. and D.G. Hardie, AMPK: Sensing Glucose as well as Cellular Energy Status. Cell Metab, 2018. 27(2): p. 299–313.

9. Han, Y., et al., Effect of metformin on all-cause and cardiovascular mortality in patients with coronary artery diseases: a systematic review and an updated meta-analysis. Cardiovascular Diabetology, 2019. 18(1): p. 96.

10. Chien, L.N., et al., Association Between Stroke Risk and Metformin Use in Hemodialysis Patients With Diabetes Mellitus: A Nested Case–Control Study. Journal of the American Heart Association, 2017. 6(11): p. e007611.

11. Dianov, G., et al., Repair pathways for processing of 8-oxoguanine in DNA by mammalian cell extracts. J Biol Chem, 1998. 273(50): p. 33811–6.

12. Wang, R., et al., OGG1-initiated base excision repair exacerbates oxidative stress-induced parthanatos. Cell Death Dis, 2018. 9(6): p. 628.

13. Zhou, X., et al., OGG1 is essential in oxidative stress induced DNA demethylation. Cell Signal, 2016. 28(9): p. 1163–71.

14. Zuo, G.F., et al., Activation of the PP2A catalytic subunit by ivabradine attenuates the development of diabetic cardiomyopathy. J Mol Cell Cardiol, 2019. 130: p. 170–183.

15. Xie, X., et al., High glucose induced endothelial cell reactive oxygen species via OGG1/PKC/NADPH oxidase pathway. Life Sci, 2020. 256: p. 117886.

16. Chao, Y., et al., Low shear stress induces endothelial reactive oxygen species via the AT1R/eNOS/NO pathway. J Cell Physiol, 2018. 233(2): p. 1384–1395.

17. Liu, Z.J., et al., OGG1 Involvement in High Glucose-Mediated Enhancement of Bupivacaine-Induced Oxidative DNA Damage in SH-SY5Y Cells. Oxid Med Cell Longev, 2015. 2015: p. 683197.

18. Zhao, Y., et al., ROS signaling under metabolic stress: cross-talk between AMPK and AKT pathway. Mol Cancer, 2017. 16(1): p. 79.

19. Hentze, M.W., et al., A brave new world of RNA-binding proteins. Nature Reviews Molecular Cell Biology, 2018. 19(5): p. 327–341.

20. Zhao, Y., et al., The lncRNA MACC1-AS1 promotes gastric cancer cell metabolic plasticity via AMPK/Lin28 mediated mRNA stability of MACC1. Mol Cancer, 2018. 17(1): p. 69.

21. Pendyala, S., et al., Regulation of NADPH oxidase in vascular endothelium: the role of phospholipases, protein kinases, and cytoskeletal proteins. Antioxid Redox Signal, 2009. 11(4): p. 841–60.

22. Ba, X., et al., 8-oxoguanine DNA glycosylase-1 augments proinflammatory gene expression by facilitating the recruitment of site-specific transcription factors. J Immunol, 2014. 192(5): p. 2384–94.

23. Griffin, S.J., J.K. Leaver, and G.J. Irving, Impact of metformin on cardiovascular disease: a meta-analysis of randomised trials among people with type 2 diabetes. Diabetologia, 2017. 60(9): p. 1620–1629.

24. Rena, G. and C.C. Lang, Repurposing Metformin for Cardiovascular Disease. Circulation, 2018. 137(5): p. 422–424.

25. Verma, S., et al., Metformin treatment corrects vascular insulin resistance in hypertension. J Hypertens, 2000. 18(10): p. 1445–50.

26. Katakam, P.V., et al., Metformin improves vascular function in insulin-resistant rats. Hypertension, 2000. 35(1 Pt 1): p. 108–12.

27. Chen, C., et al., Metformin prevents vascular damage in hypertension through the AMPK/ER stress pathway. Hypertens Res, 2019. 42(7): p. 960–969.

28. Rena, G., D.G. Hardie, and E.R. Pearson, The mechanisms of action of metformin. Diabetologia, 2017. 60(9): p. 1577–1585.

29. Davis, B.J., et al., Activation of the AMP-activated kinase by antidiabetes drug metformin stimulates nitric oxide synthesis in vivo by promoting the association of heat shock protein 90 and endothelial nitric oxide synthase. Diabetes, 2006. 55(2): p. 496–505.

30. Brynildsen, J.K., et al., Activation of AMPK by metformin improves withdrawal signs precipitated by nicotine withdrawal. Proc Natl Acad Sci U S A, 2018. 115(16): p. 4282–4287.

31. Chen, S.C., et al., Metformin suppresses adipogenesis through both AMP-activated protein kinase (AMPK)-dependent and AMPK-independent mechanisms. Mol Cell Endocrinol, 2017. 440: p. 57–68.

32. Deschemin, J.-C., et al., AMPK is not required for the effect of metformin on the inhibition of BMP6-induced hepcidin gene expression in hepatocytes. Scientific Reports, 2017. 7(1): p. 12679.

33. Tumurkhuu, G., et al., Ogg1-Dependent DNA Repair Regulates NLRP3 Inflammasome and Prevents Atherosclerosis. Circ Res, 2016. 119(6): p. e76–90.

34. Habib, S.L., et al., Tuberin regulates the DNA repair enzyme OGG1. Am J Physiol Renal Physiol, 2008. 294(1): p. F281–90.

35. Kim, H.S., et al., Potential role of 8-oxoguanine DNA glycosylase 1 as a STAT1 coactivator in endotoxin-induced inflammatory response. Free Radic Biol Med, 2016. 93: p. 12–22.

36. Fantini, D., et al., Rapid inactivation and proteasome-mediated degradation of OGG1 contribute to the synergistic effect of hyperthermia on genotoxic treatments. DNA Repair (Amst), 2013. 12(3): p. 227–37.

37. Zhang, Y., et al., NADPH oxidases and oxidase crosstalk in cardiovascular diseases: novel therapeutic targets. Nat Rev Cardiol, 2020. 17(3): p. 170–194.

38. Gorin, Y. and K. Block, Nox as a target for diabetic complications. Clin Sci (Lond), 2013. 125(8): p. 361–82.

39. Gray, S.P., et al., NADPH oxidase 1 plays a key role in diabetes mellitus-accelerated atherosclerosis. Circulation, 2013. 127(18): p. 1888–902.

40. Wang, C., et al., Diabetes aggravates myocardial ischaemia reperfusion injury via activating Nox2-related programmed cell death in an AMPK-dependent manner. J Cell Mol Med, 2020. 24(12): p. 6670–6679.

41. Williams, C.R., et al., Rosiglitazone attenuates NF-κB-mediated Nox4 upregulation in hyperglycemia-activated endothelial cells. Am J Physiol Cell Physiol, 2012. 303(2): p. C213–23.

